# Calcium binding protein Ncs1 is calcineurin-regulated in *Cryptococcus neoformans* and essential for cell division and virulence

**DOI:** 10.1101/2020.07.23.218974

**Authors:** Eamim Daidrê Squizani, Júlia Catarina Vieira Reuwsaat, Sophie Lev, Heryk Motta, Julia Sperotto, Keren Kaufman-Francis, Desmarini Desmarini, Marilene Henning Vainstein, Charley Christian Staats, Julianne T. Djordjevic, Lívia Kmetzsch

## Abstract

Intracellular calcium (Ca^2+^) is crucial for signal transduction in *Cryptococcus neoformans*, the major cause of fatal fungal meningitis. The calcineurin pathway is the only Ca^2+^-requiring signalling cascade implicated in cryptococcal stress adaptation and virulence, with Ca^2+^-binding mediated by the EF-hand domains of the Ca^2+^ sensor protein calmodulin. In this study, we identified the cryptococcal ortholog of neuronal calcium sensor-1 (Ncs1) as a member of the EF-hand superfamily. We demonstrated that Ncs1 has a role in Ca^2+^ homeostasis under stress and non-stress conditions, as the *ncs1Δ* mutant is sensitive to a high Ca^2+^ concentration and has an elevated basal Ca^2+^ level that correlates with increased expression of the Ca^2+^ transporter genes, *CCH1* and *MID1*. Furthermore, *NCS1* expression is induced by Ca^2+^, with the Ncs1 protein adopting a punctate subcellular distribution. We also demonstrate that, in contrast to *Saccharomyces cerevisiae*, *NCS1* expression in *C. neoformans* is regulated by the calcineurin pathway via the transcription factor Crz1, as *NCS1* expression is reduced by FK506 treatment and *CRZ1* deletion. Moreover, the *ncs1Δ* mutant shares a high temperature and high Ca^2+^ sensitivity phenotype with the calcineurin and calmodulin mutants (*cna1*Δ and *cam1Δ*) and the *NCS1* promoter contains two calcineurin/Crz1-dependent response elements (CDRE1). Ncs1-deficency coincided with reduced growth, characterized by delayed bud emergence and aberrant cell division, and hypovirulence in a mouse infection model. In summary, our data shows that Ncs1 plays distinct roles in Ca^2+^ sensing in *C. neoformans* despite widespread functional conservation of Ncs1 and other regulators of Ca^2+^ homeostasis.

**Importance:** *Cryptococcus neoformans* is the major cause of fungal meningitis in HIV infected patients. Several studies have highlighted the important contribution of Ca^2+^ signalling and homeostasis to the virulence of *C. neoformans*. Here, we identify the cryptococcal ortholog of neuronal calcium sensor-1 (Ncs1) and demonstrate its role in Ca^2+^ homeostasis, bud emergence, cell cycle progression and virulence. We also show that Ncs1 function is regulated by the calcineurin/Crz1 signalling cascade. Our work provides evidence of a link between Ca^2+^ homeostasis and cell cycle progression in *C. neoformans*.

## Introduction

*Cryptococcus neoformans* is a basidiomycetous pathogenic yeast, found mostly in soil and bird droppings (1–3). This pathogen is the etiological agent of cryptococcosis, which affects mainly immunocompromised individuals. This disease affects more than 220,000 HIV-infected patients per year, resulting in more than 180,000 deaths worldwide (3,4). The lung infection is initiated following the inhalation of small desiccated cells or spores. The infection can then spread via the bloodstream to the central nervous system, causing meningoencephalitis, which is the primary cause of death (1,5). To survive within the host environment, *C. neoformans* produces several virulence determinants, including a polysaccharide capsule, the pigment melanin, secreted enzymes (6–9) and extracellular vesicles (10). *C. neoformans* survival in the host is only possible due to its ability to grow at 37 °C and also aided by its capacity to survive within phagocytic mammalian cells (1,11–15).

Fungal fitness and survival in the host environment is controlled by numerous signaling pathways including those that are regulated by intracellular Ca^2+^, which is an essential second messenger in eukaryotic cells (16–19). An increase in cytosolic Ca^2+^ is monitored by Ca^2+^ sensor proteins that, upon binding to Ca^2+^, change their conformation and transduce signals onto downstream targets (20,21). An important Ca^2+^ sensor in fungal cells is calmodulin, which is a component of the calcineurin signaling pathway. Ca^2+^-induced conformational change in calmodulin activates the serine-threonine phosphatase, calcineurin. Calcineurin then mediates the regulation of several cellular responses by initiating changes in the phosphorylation status of its downstream targets (18,22,23). A major target of cryptococcal calcineurin is the transcription factor Crz1, which regulates the expression of genes involved in stress response and in the maintenance of cell wall integrity (24,25). In *C. neoformans*, the calcineurin pathway is also essential for growth at 37 °C, sexual reproduction and virulence (26); in *Saccharomyces cerevisiae*, it is required for cell cycle progression (27).

Given that high levels of cellular Ca^2+^ can be toxic, Ca^2+^ homeostasis is strictly regulated by several proteins acting as transporters, channels or pumps (28). In *C. neoformans*, these proteins include Cch1, a Ca^2+^ voltage-gated channel essential for virulence and Mid1, a stretch-activated Ca^2+^-channel, both found in the plasma membrane (19,29). Other cryptococcal calcium transporters that also promote virulence include the Ca^2+-^ATPase, EcaI, found in sarcoplasmic/endoplasmic reticulum, the H^+^/ Ca^2+^ exchanger protein Vcx1 and the Ca^2+^ ATPase Pmc1, both localized on vacuolar membranes and responsible for Ca^2+^ storage (30–33). Pmc1 is also required for *C. neoformans* transmigration through the blood-brain barrier (BBB), proving that Pmc1-regulated Ca^2+^ homeostasis is crucial for disease progression (33).

Despite the importance of Ca^2+^ homeostasis-related proteins in fungal cell fitness and virulence, with the exception of calmodulin, little is known about the function of other calcium binding proteins (CBPs) that act as Ca^2+^ sensors in *C. neoformans*. One such protein is the neuronal calcium sensor 1 (Ncs1). Ncs-1 orthologs in other fungi have roles in cell growth and viability, tolerance to Ca^2+^ (34–41), membrane sterol distribution and expression of Ca^2+^ transporter genes (41). Here, we identify and characterize the Ncs1 ortholog in *C. neoformans*. Using gene deletion and *in silico* analysis, we investigate the role of Ncs1 in Ca^2+^ homeostasis, growth, stress tolerance and virulence, and whether Ncs1 function is linked to the calcineurin pathway.

## Results

### Identification of the neuronal calcium sensor 1 (Ncs1) ortholog in *C. neoformans*

CBPs are either predominantly intrinsic membrane proteins that transport Ca^2+^ through membranes or Ca^2+^ modulated proteins, mainly represented by Ca^2+^ sensors involved in signal transduction (21,28). The later includes calmodulin and calcineurin, which harbor the calcium-binding (EF-hand) domain. Both proteins have been well studied in eukaryotic cells, including *C. neoformans* (25,26,42). However, calmodulin is the only intracellular calcium sensor characterized so far in *C. neoformans*. Considering the number and complexity of processes regulated by Ca^2+^, we sought to identify other CBPs in the EF-hand superfamily with Ca^2+^ sensor functions. In this context, we searched for the Ncs1 homologue in *C. neoformans*, given that this protein is important for Ca^2+^ regulated processes in a variety of eukaryotic cells (43).

For this purpose, we performed an *in silico* analysis at FungiDB to identify the *NCS1* coding sequence in the *C. neoformans* H99 genome (accession number CNAG_03370). Ncs-1 is well conserved in eukaryotes, with orthologs sharing common regions, such as EF hand domains and a myristoylation motif. Our analysis revealed that *C. neoformans* Ncs1 contains four EF-hands domains that span the full length of the protein (Fig. 1). Moreover, the presence of an N-terminal myristoylation motif was identified using the NMT-themyr predictor database. Myristoylation, a lipid modification conserved among eukaryotic Ncs-1 proteins (Burgoyne, 2004), is important for cell signaling, protein–protein interaction and protein targeting to endomembrane systems and the plasma membrane (45). Comparative analysis of the *C. neoformans* Ncs1 protein sequence with *Aspergillus fumigatus NCSA* (Afu6g14240), *Schizossacharomyces pombe NCS1* (SPAC18B11.04) and *S. cerevisiae FRQ1* (YDR373W), which are already functionally characterized (34,40,41), revealed a high amino acid sequence similarity (86, 87 and 81%, respectively) (Fig. 1).

**Figure 1.**
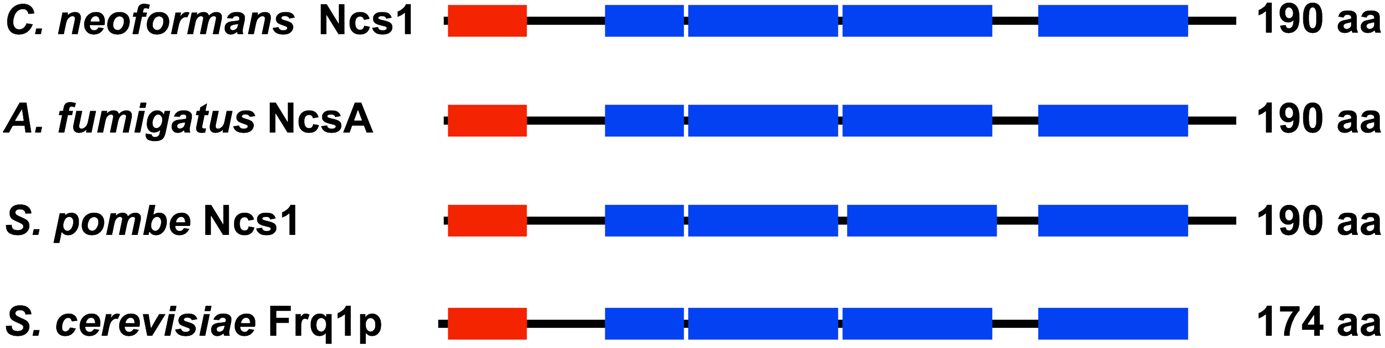
Identification of Ncs1 as a putative calcium binding protein in *C. neoformans*. Comparative *in silico* analysis of *C. neoformans* Ncs1 (CNAG_03370), *Aspergillus fumigatus* NcsA (Afu6g14240), *Schizossacharomyces pombe* Ncs1 (SPAC18B11.04) and *S. cerevisiae* Frq1 (YDR373W) amino acid sequences indicates the presence and position of the four EF-hand domains (blue bars) and the N-terminal myristoylation domain (red bars).

### Disruption of the *NCS1* gene affects *C. neoformans* traits associated with calcium homeostasis

Calcium sensor proteins measure fluctuations in free cytosolic Ca^2+^ and transduce the signal to downstream effectors (41,42,46). To determine whether Ncs1 plays a similar role in *C. neoformans*, we obtained a *NCS1* gene knockout strain (*ncs1Δ*) from the Madhani’s mutant collection and generated a *NCS1* reconstituted (*ncs1Δ::NCS1*) strain (Fig. S1) using the *ncs1Δ* background. We then evaluated the ability of these mutant strains to grow under different stresses. We initially chose high Ca^2+^ concentration (to alter Ca^2+^ homeostasis) and high temperatures (37 °C and 39 °C), as the calcineurin *(cna1Δ)* and calmodulin *(cam1Δ)* mutants were shown to be sensitive under these growth conditions (24,25,42,42). We observed impaired *ncs1Δ* growth in high Ca^2+^ levels and at 39°C, but not at 37°C; these growth defects were restored to WT levels in the *ncs1Δ::NCS1* strain (Fig. 2). Other traits associated with Ca^2+^-calcineurin pathway, such as growth in the presence of cell wall perturbing agents (Calcofluor white and Congo red) and osmotic stress (1 M NaCl) were evaluated in the *ncs1*Δ null mutant, with no effect observed (Fig S2).

**Figure 2.**
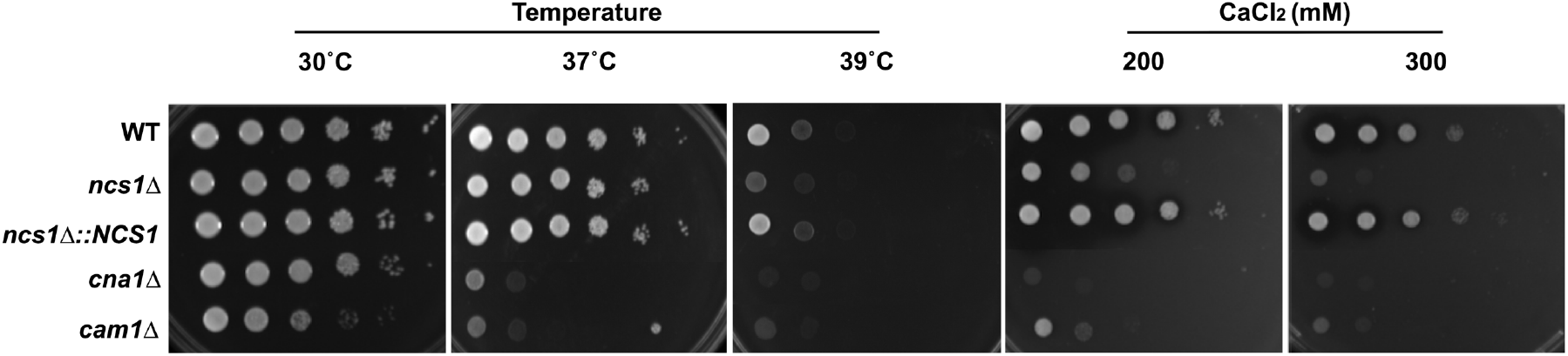
Disruption of *NCS1* leads to stress sensitivity in *C. neoformans*. Spot plate assays of the WT, *ncs1Δ* mutant and *ncs1Δ::NCS1* complemented cells were performed on YPD agar. The plates were incubated at 30 °C (control for normal growth), 37 °C or 39 °C, and under stress induced by Ca^2+^ (200 mM or 300mM CaCl_2_, at 30 °C). The calcineurin (*cna1*Δ) and calmodulin (*cam1Δ*) mutants were included as controls. All assays were conducted for 48 h.

We also evaluated whether the level of free intracellular Ca^2+^ in *C. neoformans* is affected in the absence of Ncs1. Relative to the WT strain, the *ncs1Δ* mutant had a higher basal level of free cytosolic Ca^2+^, which was reduced to WT levels in the *ncs1Δ::NCS1* strain. This high Ca^2+^ level phenotype was shared with that observed for the *cna1Δ* and *cam1Δ* mutant strains (Fig 3A). In *S. pombe*, Ncs1 physically interacts with the Mid1 ortholog, Yam8, which is a stretch-activated Ca^2+^-channel. *S. pombe YAM8* gene disruption in the *ncs1Δ* background restored the Ca^2+^ sensitive phenotype (35). In this study, the authors proposed that Ncs1 and Yam8 ortholog cooperate to maintain intracellular Ca^2+^ homeostasis. We therefore investigated whether Mid1 and another plasma membrane Ca^2+^ transporter, Cch1, are responsible for the increased intracellular Ca^2+^ observed in the *ncs1Δ* mutant. Specifically, we used real-time RT-qPCR to compare the expression of *MID1* and *CCH1* in WT and *ncs1Δ* grown in the absence or presence of Ca^2+^ (100 mM CaCl_2_ for 24 hours). No differences in *CCH1* and *MID1* expression levels were observed in WT and *ncs1Δ* grown in the absence of Ca^2+^ (Fig 3B). However, expression of both genes increased by approximately 3-fold in the *ncs1Δ* mutant strain following growth in the presence of Ca^2+^ (Fig 3B). This suggests that Cch1 and Mid1 could be the potential source of the extra Ca^2+^ in the *ncs1Δ* mutant, since they import Ca^2+^ to the cytosol (29,47). We also generated a *mid1Δncs1Δ* double mutant in *C. neoformans* to evaluate if calcium sensitivity would be restored. However, increased intracellular Ca^2+^ accumulation and Ca^2+^ sensitivity persisted in the *mid1Δncs1Δ* double mutant (Fig. S3), suggesting that Mid1 and Cch1 are functionally redundant.

**Figure 3.**
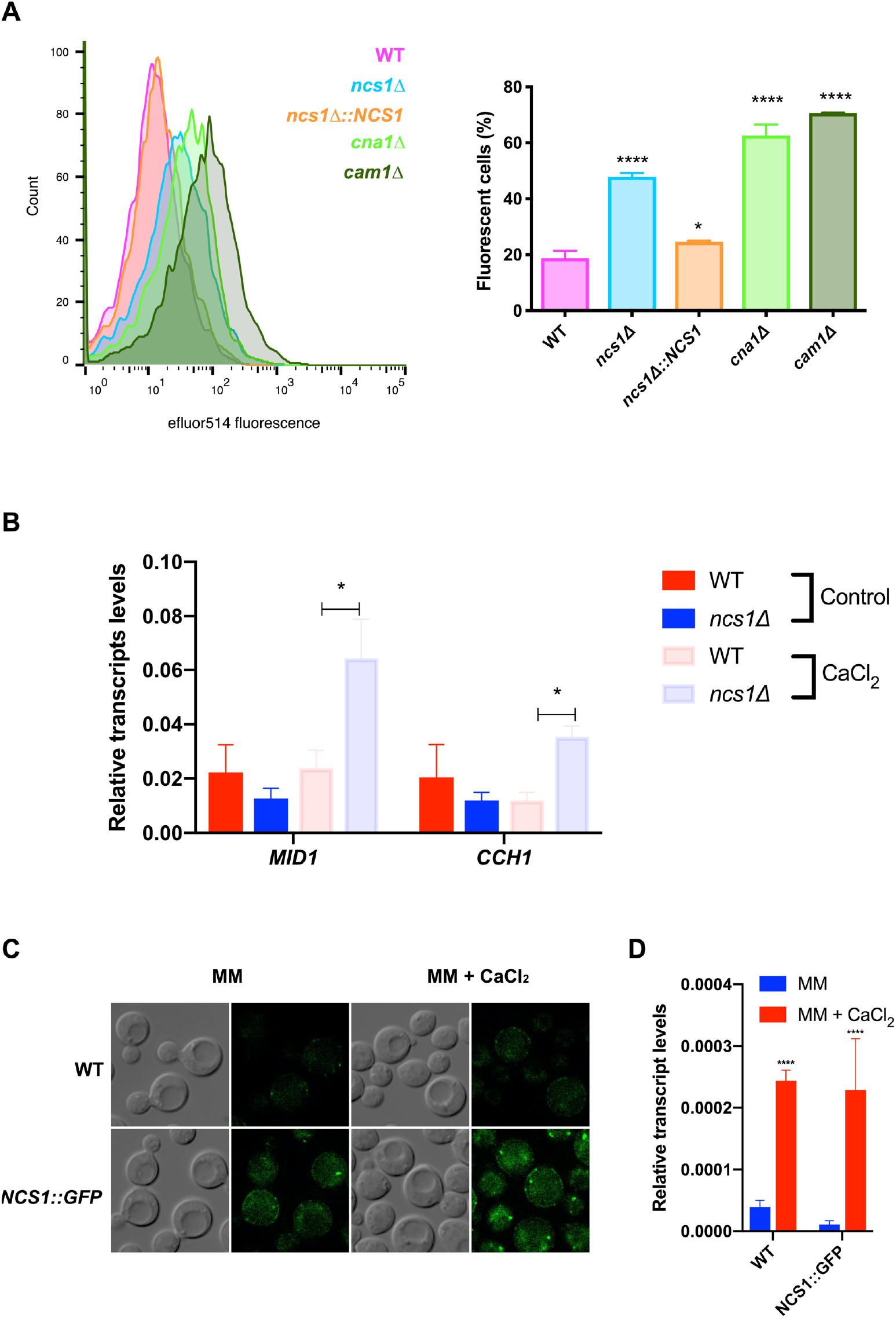
Cryptococcal Ncs1 is associated with Ca^2+^ homeostasis. (A) The basal levels of free intracellular Ca^2+^ in WT, *ncs1Δ, ncs1Δ::NCS1, cna1Δ, and cam1Δ* mutant cells were quantified by flow cytometry following staining with the calcium specific dye, Fluo-4-AM. Left panel represents the histogram of Fluo-4-AM emitted fluorescence of indicated strains cultivated in YPD medium at 30 °C. Right panel represents the percentage of gated fluorescent cells ± standard deviation (three biological replicates). Mean values were compared using one-way-ANOVA and Dunnet’s as a *post hoc* test. Statistical significance is represented as **** *p*< 0.0001 and * *p*< 0.05. (B) The transcript levels of genes encoding the calcium transporters, *CCH1* and *MID1*, were evaluated using RT-qPCR. The WT and *ncs1Δ* strains (10^6^ cells/mL) were incubated in YPD for 16 h with shaking, either at 37 °C (control), or 37 °C supplemented with Ca^2+^ (100 mM CaCl_2_). RNA was extracted and cDNA synthesized. Each bar represents the mean ± the standard deviation (n=3) for each gene in each strain normalized to actin. Statistical analysis was performed using a *Student’s t test* (* *p*< 0.05). (C) Ncs1 was tagged with GFP (*NCS1::GFP)* and the effect of CaCl_2_ supplementation on Ncs1 production and subcellular localization was assessed by fluorescence microscopy. YPD overnight cultures of WT (autofluorescence background control) and the *NCS1::GFP* strain were washed twice with water and used to seed on minimal media (MM) or MM supplemented with Ca^2+^ (100mM CaCl_2_) at OD_600nm_ = 1. The cultures were further incubated for 4 hours at 30°C prior to visualization. DIC and green fluorescent images were included. (D) The cultures prepared in (C) were also used to extract RNA and perform RT-qPCR to assess the effect of Ca^2+^ on the transcript levels of *NCS1*, normalized to actin. Statistical analysis was performed using one-way ANOVA with Tukey’s *post-hoc* test. Comparisons were conducted between WT cells grown in the absence or in the presence of Ca^2+^ or between *NCS1::GFP* cells grown in the absence or in the presence of Ca^2+^. Statistical significance is indicated as *p \*\*\*\** < 0.0001.

We also C-terminally tagged Ncs1 with GFP (*NCS1::GFP*) to assess Ncs1 subcellular localization. Faint Ncs1 fluorescence was observed when the strain was cultured in the absence of Ca^2+^, however it was higher than that observed for the non-fluorescent WT control strain (Fig. 3C). The fluorescent strain cultured in the presence of Ca^2+^ (100mM CaCl_2_) produced a more intense signal and Ncs1 adopted a more punctuate staining pattern: 24.5 ± 1.1 % and 36.3 ± 3.0 % of the cell population displayed puncta both in the absence (MM) and presence of Ca^2+^ (MM + CaCl_2_), respectively (*p* =0.013 Welch’s test with n ≥ 200 cells per sample) (Fig. 3C). Increased Ncs1 fluorescence in the presence of Ca^2+^ correlated with higher expression of *NCS1* by *NCS1::GFP* strain in the same condition (Fig. 3D). Taken together, these results suggest that Ncs1 responds to increase in intracellular Ca^2+^ levels and participates in the regulation of calcium homeostasis in *C. neoformans*.

### *NCS1* is a calcineurin-Crz1 responsive gene

Given that *NCS1* is a Ca^2+^-responsive gene in *C. neoformans* (Fig. 3D), we investigated whether *NCS1* expression is regulated by the calcineurin signaling pathway via the transcription factor Crz1. *NCS1* expression was analyzed in the presence and absence of the calcineurin inhibitor FK506 (Fig. 4A), and in the WT and *crz1*Δ mutant (Fig. 4B). The results demonstrated that FK506-treatment reduced *NCS1* transcription in the WT (Fig 4A), and that *NCS1* expression was downregulated in the *crz1*Δ mutant at 25 °C and 37 °C (Fig. 4B). In further support of *NCS1* being a Crz1 target, we identified two Crz1-binding consensus motifs (48) in the putative *NCS1* regulatory region encompassing the 1,000 nucleotides sequence upstream of the transcription start site (Fig. 4C). These findings provide evidence that Ncs1 and calcineurin work together to regulate Ca^2+^ homeostasis.

**Figure 4.**
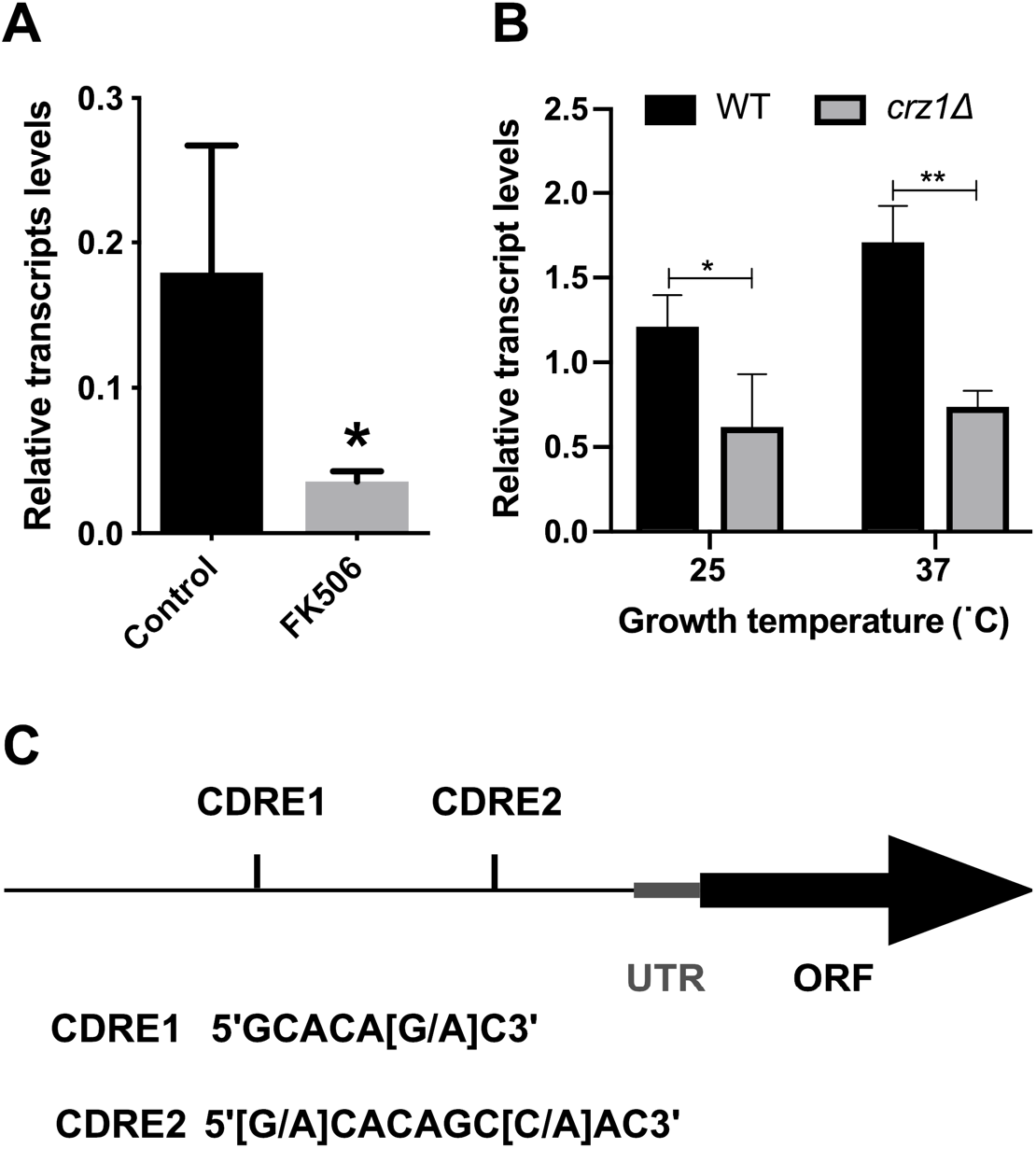
*NCS1* gene expression is regulated by Crz1. (A) The transcript levels of *NCS1* were determined in conditions of calcineurin inhibition. Yeast cells were incubated in YPD media at 37 °C in the absence or presence of FK506 (1 μg/mL) for 1 h. *NCS1* expression was normalized to *ACT1* transcript levels. Bars represent the mean ± standard deviation (three biological replicates). Statistics were conducted using *Student’s t-test*. (\**p* < 0.05, *** *p* < 0.001). (B) *NCS1* gene expression in WT and *crz1Δ* null mutant cells were assessed by RT-qPCR. Yeast cells were incubated in YPD at 25 °C or 37 °C, for 16 h. *NCS1* expression was normalized to *ACT1* transcript levels. Each bar represents the mean ± the standard deviation (three biological triplicates). Statistical analysis was performed using *Student’s t-test* (\**p* < 0.05 and ** *p* < 0.01). (C) The *NCS1* regulatory sequence contains two Crz1 binding motifs (CDRE1 and CDRE2). CDRE, calcineurin dependent response element, UTR, untranslated region, ORF, open reading frame.

### Ncs1 activity is essential for *C. neoformans* virulence

As proven in other studies, the disruption of non-Ncs1-mediated Ca^2+^ homeostasis mechanisms is important for cryptococcal pathogenicity (29–33,49). To determine whether disruption of Ncs1-mediated calcium homeostasis also contributes to pathogenicity, we compared the virulence of the *ncs1Δ* mutant strain to that observed for the WT and *ncs1Δ::NCS1* strains in a mouse inhalation model of cryptococcosis. In a Kaplan-Meier survival study, the *ncs1Δ* null mutant strain was found to be hypovirulent (Median Lethal Time-LT_50_ 32.7 days) when compared to WT (LT_50_ 18.9, p < 0.0001) and the *ncs1Δ::NCS1* strains (LT_50_ 17.4 days, p < 0.0001) (Fig. 5A). Although the disruption of *NCS1* prolonged mouse survival, no difference in the fungal burdens in lung and brain were observed at time of death when infected mice had lost 20% of their pre-infection weight (Fig. 5B). Thus the *ncs1Δ* null mutant strain is capable of infecting the lung and brain tissue but potentially grows at a slower rate compared to WT and the *ncs1Δ::NCS1* strains.

**Figure 5.**
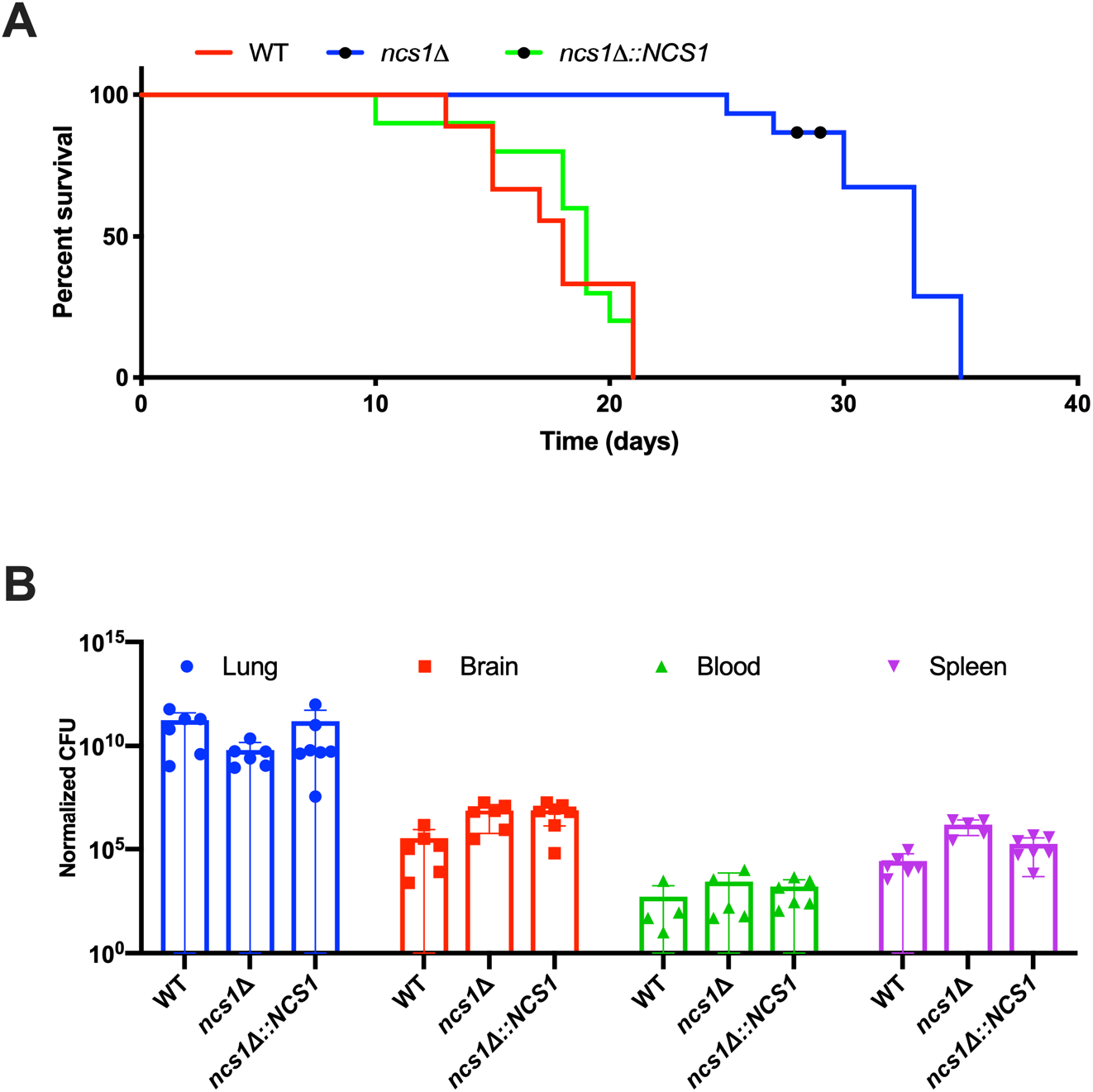
Ncs1 is required for full virulence in a mouse inhalation model of cryptococcosis. C57BL/6J mice (10 mice per group) were infected with 500,000 cells of WT, *ncs1Δ* or *ncs1Δ::NCS1* strains. Mice were monitored daily and euthanized by CO_2_ asphyxiation when they had lost 20% of their pre-infection weight. In (A), median mouse survival differences were estimated using a Kaplan-Meier Log-rank Mantel-Cox test. The increase in median survival of *ncs1Δ-*infected mice relative to the other two infection groups was statistically significant (*p* < 0.0001). In (B) lungs, brain and spleen were removed post euthanasia, weighed, homogenized, serially diluted and plated onto Sabouraud dextrose agar plates to determine fungal burden by quantitative culture (CFUs) following 3 days growth at 30°C. CFUs were adjusted to reflect CFU/gram of tissue and CFU/mL blood (normalized CFU). Statistical significance was determined using one-way ANOVA. However, no differences in organ burden were found.

### Ncs1 is necessary for growth under host mimicking conditions

We also analyzed the capability of *ncs1Δ* mutant to synthesize the polysaccharide capsule, since this is the main cryptococcal virulence factor (1,12). We observed that when the *ncs1Δ* strain is grown under capsule inducing conditions (DMEM at 37 °C and 5 % CO_2_), mutant cells produce smaller capsules in comparison to WT and *ncs1Δ::NCS1* strain (Fig. 6A). However, capsule size was not affected following growth in mouse serum (data not shown). Next, we compared growth of the *Δncs1* mutant to that of the WT and *ncs1Δ::NCS1* strains under conditions that more closely mimic those found in the mammalian host (DMEM at 37 °C and 5 % CO_2_). We found that the mutant growth was drastically compromised (Fig. 6B). Similarly, growth of *ncs1Δ* was severely impaired in mouse serum over a 24-hour period at 37 °C with 5% CO_2_ (Fig. 6C). Collectively, these results suggest that hypovirulence of the *ncs1Δ* mutant is most likely associated with the observed growth defects, rather than the reduced capsule size phenotype detected upon cells cultured in DMEM.

**Figure 6.**
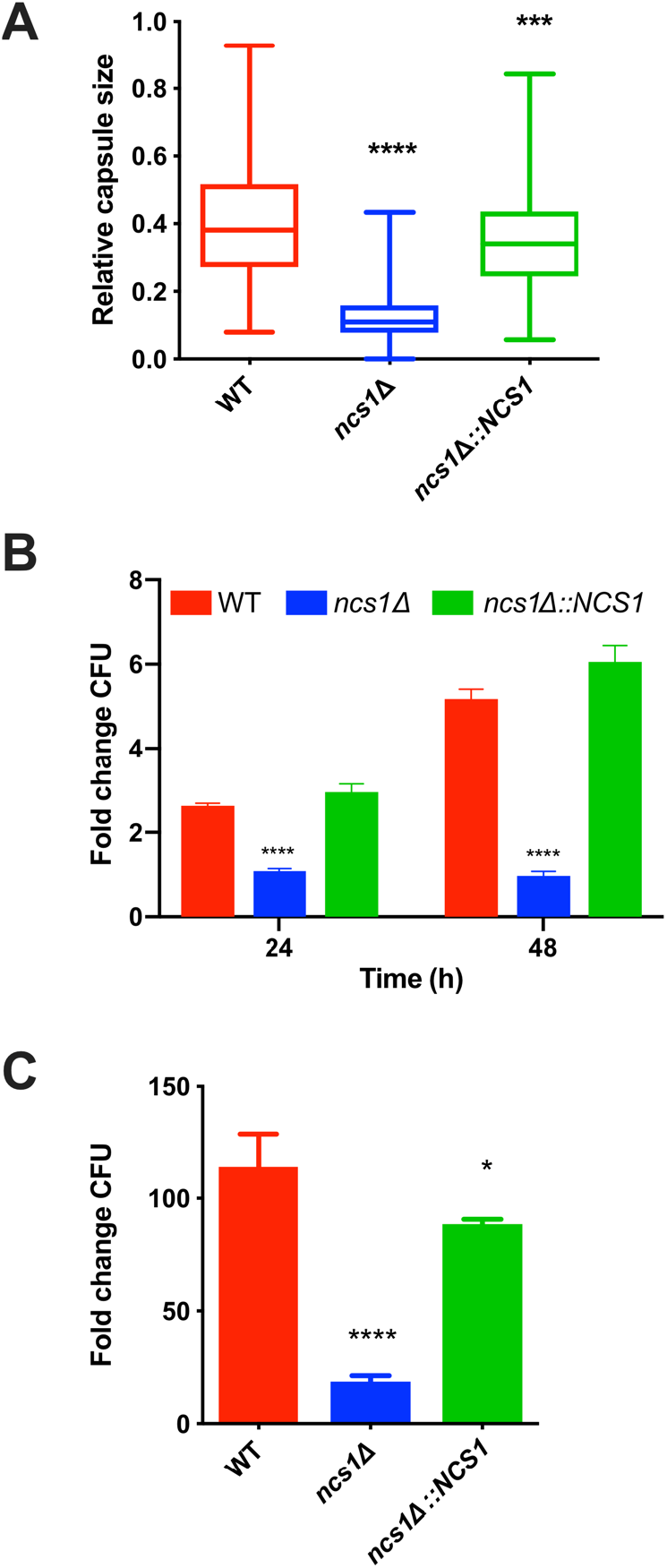
Ncs1 is necessary for growth under host mimicking conditions. (A) Capsule size of WT, *ncs1Δ* and *ncs1Δ::NCS1* cells was determined following incubation in capsule inducing media (DMEM) for 72 h (37 °C 5% CO_2_). Capsules were visualized by India ink staining and light microscopy, and measurements were performed using Image J software for at least 50 cells of each strain. Statistical analysis was performed using one-way ANOVA, with Tukey *post-hoc* test. Statistical significance is represented as **** *p* < 0.0001 and *** *p* < 0.001, as compared to the WT. (B) Growth of the WT, *ncs1Δ* and *ncs1Δ::NCS1* cells in DMEM (37 °C 5% CO_2_) for 24 or 48 h was assessed by quantitative culture (CFU). The results represent the mean ± standard deviation (three biological replicates) of each strain normalized to the CFU of the inoculum, described as fold change. Statistical analysis was performed using one-way ANOVA with Dunnet’s *post-hoc* test. Significant differences compared to WT are marked (**** *p* < 0.0001). (C) Growth of the WT, *ncs1Δ* and *ncs1Δ::NCS1* cells for 24 h at 37 °C 5% CO_2_ in heat-inactivated mouse serum was indicated by quantitative culture (CFU). The results are expressed as a fold change relative to the initial inoculum (10^4^ cells/mL), and represent the mean ± standard deviation (three biological replicates). Statistical analysis was performed using one-way ANOVA and Dunnet’s *post –hoc* test (* *p* < 0.05 and **** *p* < 0.0001 relative to WT).

### Ncs1 is important for the release of daughter cells

Microscopic analysis to evaluate the size of the polysaccharide capsule and the growth rate in mouse serum revealed that some *ncs1Δ* cells displayed aberrant morphology and cell division (Fig. 7A), suggesting that Ncs1 could play a role in cell cycle progression. We therefore investigated the growth defect further by determining the time it took for buds to emerge using time-lapse microscopy (Fig. 7B and Movies S1 and S2). Given that the mutant was severely attenuated in growth when cultured in DMEM or exposed to mouse serum, we chose YPD medium for this analysis, as it is a richer medium where mutant growth is not as compromised. To avoid bias due to lack of synchronization, we only measured the time of bud emergence in cells after the bud of the first daughter cell had separated from the mother or, in the case of the mutant cells, where progeny did not detach from mother cell, after the second bud emergence. The results demonstrate that it took ~70 min for buds to emerge in the WT cells and more than 140 min for buds to emerge in isolated and clumped *ncs1*Δ mutant cells (Fig. 7C). Furthermore, buds were slow to be released in some *ncs1*Δ mutant cells, resulting in more extensive cell clumping.

**Figure 7.**
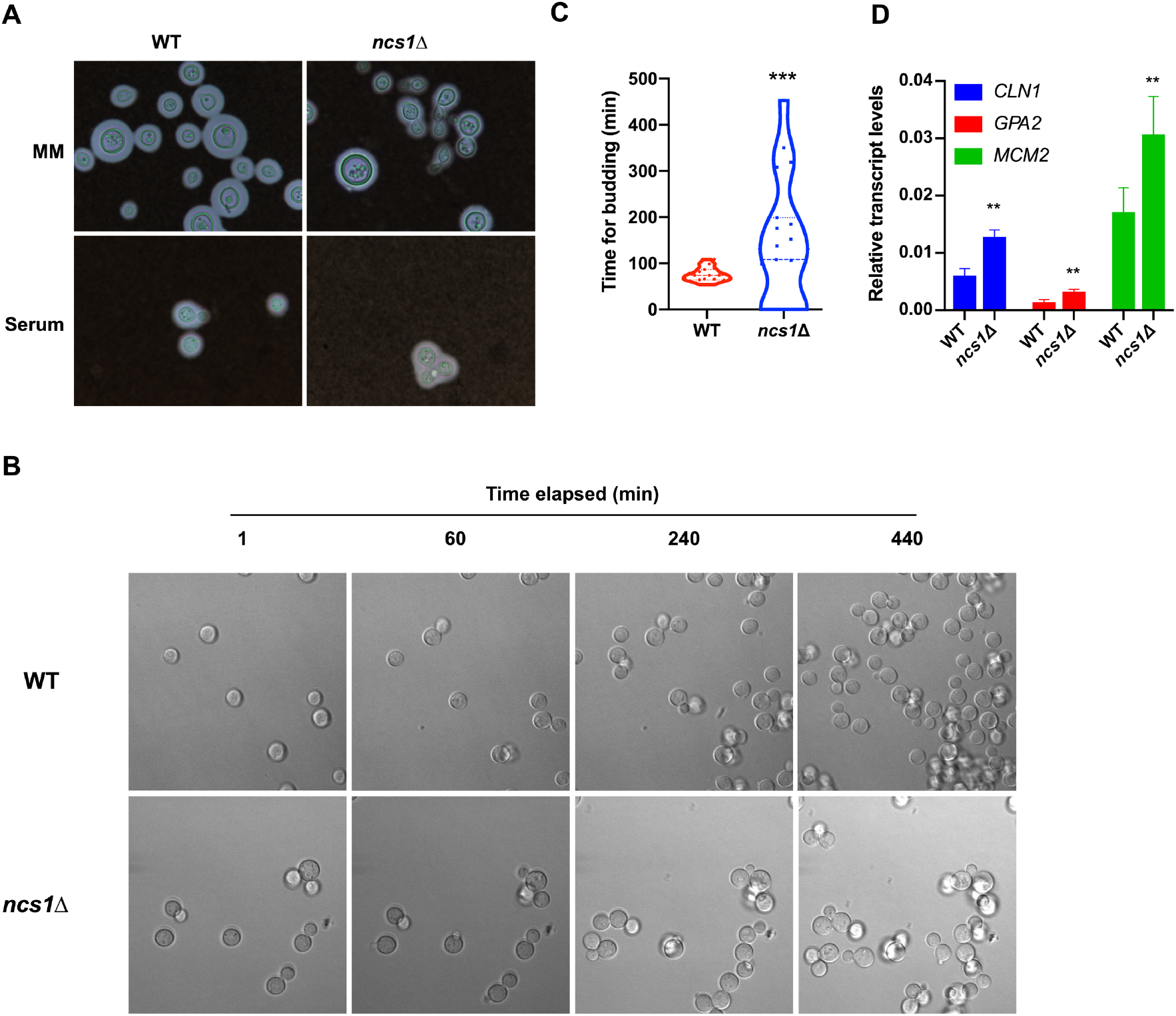
The *ncs1Δ* null mutant strain displays aberrant cell division and morphology (A), delayed bud emergence (B, C) and altered cell cycle regulation (D). (A) WT and *ncs1Δ* cells were grown in minimal media (MM) for 72 h at 37 °C and 5% CO_2_ (upper panel) or in heat-inactivated mouse serum for 24 h at 37 °C and 5% CO_2_ (lower panel), stained with India ink and visualized by light microscopy. (B and C) Fungal cells were incubated in YPD medium for 16 h inside a chamber coupled to a confocal microscope (37 °C, 5% CO_2_), and bud emergence time was recorded using time-lapse microscopy. Time measurements were initiated after the first round of bud emergence to avoid errors associated with the lack of synchronization. Images were acquired every 30 seconds. The graph in (C) represents the mean time for buds to emerge (minutes) ± standard deviation of at least 15 cells per strain. Statistical analysis was performed using the nonparametric Mann-Whitney *test* (*** *p*< 0.0001). (D) Transcript levels of genes encoding cell cycle regulators were assessed in WT and *ncs1Δ* cells by RT-qPCR. Cells were grown in YPD at 37 °C for 4 hours. Results represent the mean transcript level ± standard deviation (three biological triplicates) with each gene normalized *ACT1* transcript levels. Statistical analysis was performed using *Student’s t-test* (** *p* < 0.01).

As cell division is linked to the cell cycle, we evaluated whether cells lacking *NCS1* displayed defects in cell cycle regulation by measuring the levels of two transcripts associated with different stages of the cell cycle: the G_1_ cyclin encoded by *CNL1* (50) and the S phase DNA replication licensing factor encoded by *MCM2* (51,52). We also measured the transcript levels of the G protein coupled receptor encoded by *GPA2*, which displays oscillatory expression during the cell cycle (51,52). All three genes were upregulated in the *ncs1Δ* strain compared to WT after 4 h of growth in YPD (Fig. 7D), reinforcing that cell cycle progression is altered in the *ncs1Δ* mutant strain.

## Discussion

Our results indicate that *NCS1* expression in *C. neoformans* is regulated by Ca^2+^ and the calcineurin/Crz1 pathway and corroborate findings on the Ncs1 ortholog in fission yeast (35). In contrast to our conclusions and those made in studies using *S. pombe*, the *S. cerevisiae* Ncs1 ortholog, Frq1, was found to be essential for viability and the level of *FRQ1* expression was not influenced by the calcineurin/Crz1 pathway as revealed by microarray analysis (53). This suggests that distinct calcium sensing mechanisms exist in fungal species despite wide-spread functional conservation of Ncs1 and other regulators of Ca^2+^ homeostasis.

An interesting feature of the *ncs1Δ* mutant is its attenuated virulence in a murine model of cryptococcosis. In contrast, attenuated virulence was not observed for the null Ncs1 ortholog mutant (*NCSA)* in *A. fumigatus* (41) reaffirming that processes regulated by Ncs1 orthologs in pathogenic fungi differ, or that other genes can compensate in the absence of *NCSA*. Moreover, cryptococcal *ncs1Δ* took longer to achieve the growth densities associated with debilitating infection in the tissues of WT-infected mice. This slower growth phenotype *in vivo* correlated with the reduced rate of proliferation of *ncs1Δ* in mouse serum and impaired bud emergence and release. These results confirm that Ncs1 is important for fungal adaptation to the host environment and for the establishment of disease and reaffirm the importance of Ca^2+^ homeostasis and Ca^2+^ signaling in cryptococcal virulence. Our findings also extend the set of calcium-related genes involved in virulence to include Ncs1.

Given that expression of ~40 virulence-associated genes is linked to the cell cycle in *C. neoformans*, the control of this process is fundamental to disease progression (51). In *S. cerevisiae*, Ca^2+^ homeostasis is linked to cell cycle regulation as a decrease in intracellular Ca^2+^ leads to transient arrest in the G_1_ phase, followed by interruption in the G_2_/M phase (27,54–56). Moreover, bud emergence and the cell cycle depend on calcineurin activity, which regulates the availability of proteins involved in cell cycle regulation. These proteins include Swe1, a negative regulator of Cdc28/Clb complex; Cln2, a protein kinase required for cell cycle progression, and a G_2_ cyclin (27). A role for calcineurin signaling in cell cycle regulation in *C. neoformans* is supported by the fact that a downstream target gene of the calcineurin pathway, *CHS6*, is periodically expressed throughout the cell cycle (24,51,52) and by our new data demonstrating that Ncs1 interferes with the transcription profile of genes associated with cell cycle progression (*CLN1, GPA2* and *MCM2)*. Notably, overexpression of the G1/S cyclin, Cln1, *S. cerevisiae*, led to a filamentation phenotype (57). In our study, a cell cycle regulation imbalance in the cryptococcal *ncs1*Δ mutant correlated with impaired bud emergence in time-lapse microscopy images of *ncs1Δ* cells. Furthermore, the *C. neoformans cln1Δ* mutant exhibited aberrant bud emergence and cell division and a consistent delay in budding (58). All of these phenotypes are shared with *ncs1Δ* mutant, reinforcing the connection between Ca^2+^ signaling and cell cycle progression in *C. neoformans*.

We hypothesize that Ca^2+^ excess, or even other types of stress, leads to activation of the calcineurin pathway, which ultimately drives the expression of Ncs1 in a Crz1-dependent fashion. Therefore, Ca^2+^-activated Ncs1 would participate in a diverse array of cellular processes to cope with Ca^2+^ excess, including the regulation of cell division via its potential association with Pik1, a protein implicated in cell septation in fission yeast (59). Two lines of evidences support this hypothesis: (i) yeast Pik1 forms puncta consistent with its localization in the Golgi apparatus (60) and we observed that cryptococcal Ncs1 also forms puncta, particularly when Ca^2+^ is present; and (ii) yeast Frq1 physically interacts with Pik1 (60). Structural studies performed on Ncs1 reveal that the myristoyl group flips out following Ca^2+^ binding, allowing Ncs1 to anchor reversibly to membranes (20,39,40). Thus, it is possible that Ca^2+^ binding to Ncs1 exposes the hydrophobic N-myristoylation domain, promoting Ncs1 association with Pik1 in Golgi membranes and, hence, proper cell septation and division.

In summary, we have characterized the Ncs1 homologue in *C. neoformans*, demonstrating its importance in Ca^2+^ homeostasis and virulence. We showed that, in contrast to *S. cerevisiae*, *NCS1* is a calcineurin-responsive gene in *C. neoformans*, with calcineurin and Ncs1 working together to regulate calcium homeostasis and, hence, promote fungal growth and virulence. To our knowledge, this is the first report of a role for Ncs1 in fungal virulence using a mammalian infection model, and of a correlation between Ca^2+^ signaling and cell cycle progression in *C. neoformans*.

## Material and methods

### Fungal strains and media

The *C. neoformans* serotype A strain Kn99 was chosen to conduct the study as wild type (WT). The *NCS1* gene (CNAG_03370) deletion mutant (*ncs1*Δ), *cna1Δ* mutant and *cam1Δ* mutant were obtained from Dr. H. Madhani’s library (61). The *ncs1*Δ reconstituted strain (*ncs1Δ::NCS1)*, the *mid1*Δ*ncs1*Δ double mutant and the *NCS1::GFP* strains were all constructed using overlapping PCR as previously described (62), and site directed homologous recombination was performed. Transformation was carried out using biolistic transformation, as previously described (63). The primer list is presented at Table S1, and the confirmations of the cassette’s insertions are demonstrated in Fig. S1. Fungal cells were maintained on solid YPD medium (1 % yeast extract, 2 % peptone, 2 % dextrose and 1.5 % agar). YPD plates containing hygromycin (200 μg/mL) or G418 (100 μg/mL) were used to select *C. neoformans* transformants.

### *In silico* analysis

To evaluate Ncs1 protein conserved domains we used the protein sequences and annotations retrieved from FungiDB database (http://fungidb.org) (64), applying FungiDB tools and InterproScan database (65). The same was performed for NMT-themyrpredictor database to identify the N-terminal myristoylation consensus sequence. Conservation was assessed using BLASTp against target proteins. Finally, the presence of the Crz1-binding motif on the promoter region of the *NCS1* gene was made by manually search. We recovered the putative regulatory regions of cryptococcal genes from FungiDB (http://fungidb.org), selecting 1,000 bp upstream of the transcription start site of *NCS1* gene. The sequences utilized for Crz1-binding motif search are already described (48).

### Virulence assay

Virulence assays were performed as previously described (66). Briefly, female C57BL/6 mice (10 per infection group) were anesthetized by inhalation of 3% isoflurane in oxygen and infected with 5 × 10^5^ fungal cells (WT, *ncs1Δ* or *ncs1Δ::NCS1* strains) via the nasal passages. Mice were monitored daily and euthanized by CO_2_ asphyxiation when they had lost 20% of their pre-infection weight, or prior in case of debilitating symptoms of infection. Median survival differences were estimated using a Kaplan-Meier Log-rank Mantel-Cox test. Post euthanasia, lungs and brain were removed, weighed, and homogenized in 2 ml sterile PBS using a BeadBug (Benchmark Scientific). Organ homogenates were serially diluted and plated onto Sabouraud dextrose agar plates. Plates were incubated at 30°C for 2 days. Colony counts were performed and adjusted to reflect the total number of colony-forming units (CFU) per gram of tissue or ml of blood. For fungal burden analysis, Two-way ANOVA with Tukey post-hoc was utilized to determine the statistical significance.

### *C. neoformans* replication in mice serum

A total of ten BALB/c mice (10-week-old) were obtained from Biotechnology Center, UFRGS, Brazil. Mice were anaesthetized using isoflurane (in a chamber), and blood was collected from mice’s retro-orbital space, using a glass capillary. Next, mice were euthanized using an overdose of thiopental (140 mg/kg). Serum was obtained from total blood after centrifugation (3000 x *g*, 15 min at room temperature). A total of 1,000 cells in 100 μL suspension of the WT, *ncs1Δ* and *ncs1Δ::NCS1* strains were inoculated at heat-inactivated mice serum in a 96-well plate and incubated at 37 °C, 5 % CO_2_ for 24 h. Thereon, yeast cells were collected and plated on YPD plates for CFU determination. Separate wells were conducted to cell morphology analysis, where yeast cells were firstly fixed with 4% paraformaldehyde for 30 minutes at 37 °C, and then analyzed using light microscopy.

### Yeast growth in DMEM

A total of 1 × 10^6^ cells in 1000 μL suspension of the WT, *ncs1Δ* and *ncs1Δ::NCS1* strains were inoculated at DMEM media in a 24-well plate and incubated at 37 °C, 5 % CO_2_ for 24 and 48 h. Next, yeast cells were gathered and plated on YPD plates for CFU determination. Separate wells were conducted to cell morphology analysis, where fungal cells were fixed with 4% paraformaldehyde for 30 minutes at 37 °C, and then analyzed using India ink counterstaining in a light microscopy.

### Intracellular calcium measurements

Free intracellular Ca^2+^ in *C. neoformans* was quantified by flow cytometry (Millipore Guava-soft) following cellular staining with the Calcium Sensor Dye Fluo-4-AM (Termofisher Scientific) at a final concentration of 2 μM. Briefly, yeast cells were cultured overnight on YPD at 30 °C with shaking. Next, cells were centrifuged (6000 rpm for 3 minutes) and washed twice with phosphate buffer. After adjusting the cell density (OD_600nm_= 1.0), the Fluo-4-AM dye was added to each tube and incubated at 37°C for 1 h. The flow was adjusted to pass < 500 cells/μL, and a total of 5,000 events were evaluated.

### Phenotypic characterization assays

For phenotypic characterization, WT, *ncs1Δ* mutant, and *ncs1Δ::NCS1* complemented strains were grown overnight on YPD at 30 °C with shaking. Further, cells were centrifuged and washed twice with deionized water, and adjusted to 10^8^ cells/mL. The cell suspensions were then subjected to serial dilution (10-fold), and 3 μL of each dilution was spotted onto YPD agar supplemented with different stressors, including CaCl_2_ (200 mM and 300mM). Cell wall perturbation was assessed using Congo red (0.1 %) and Calcofluor white (0.5 mg/ mL), as previously described (24). The sensitivity to osmotic stress was evaluated utilizing NaCl 1M. Morover, menadione (30 μM) was used as an oxidative stressor, and low phosphate environment was used as a starvation condition (67). All the plates were incubated for 48 hours at 30 °C and photographed, with the exception of plates incubated at high temperatures (37 °C or 39 °C).

### Fluorescence and light microscopy

Fluorescence microscopy assays were accomplished using a DeltaVision fluorescence microscope. WT and *NCS1::GFP* cells were incubated at minimal media (2 g/L L-asparagine, 1 g/L MgSO_4_. 7H_2_O, 6 g/L KH_2_PO_4_, 2 g/L thiamine) without or supplemented with 100 mM CaCl_2_ for 16 h at 30 °C with shaking. Thereafter, cells were washed once with PBS and analyzed. For light microscopy, WT, *ncs1Δ* mutant, and *ncs1Δ::NCS1* cells were grown in DMEM or minimal media, at 37 °C, and 5% CO_2_ for 72 h. Next, the cells were fixed with 4% paraformaldehyde for 30 min at 37 °C, washed with PBS and then analyzed under light microscopy, using counterstaining with India ink. To define the relative capsule sizes, measures of the distance between the cell wall and the capsule outer border were determined and divided by each cell diameter, through IMAGEJ software (http://rsbweb.nih.gov/ij/). At least 50 cells of each strain were measured.

### Time-lapse microscopy

Cellular division was followed using confocal microscopy. The experimental design was performed as already described (50), with few modifications. Briefly, WT or *ncs1Δ* mutant cells were cultured overnight on liquid YPD medium at 30 °C with shaking. Further, cells were washed twice with PBS and adjusted to 10^6^ cells/mL with YPD medium at pH=7.45. One hundred μL of cell suspension was inoculated on a cell culture dish, 35/10 mm glass bottom (Greiner Bio-one). The culture dish was previously treated with 100 μL of poli-L-lysin (0.1mg/mL) for 1 h, washed 3 times with PBS, and then incubated with 10 μg/mL MAb 18B7 for 1 h. Thereon, the culture dishes were incubated in a temperature-controlled microscope chamber adjusted to 37 °C, and 5 % CO_2_. Image acquisition was done in a 30-seconds interval, using differential inference contrast (DIC) objective in a confocal microscope FV1000, at Microscopy and Microanalysis Center (CMM) of the Universidade Federal do Rio Grande do Sul (UFRGS). The statistical analysis was done by timing how long each mother cell took to originate a bud. Measurements were performed at the beginning of the second budding in order to avoid errors associated with the lack of tools to synchronize cells.

### Quantitative RT-qPCR analysis

For gene expression analysis, strains were subjected to different conditions, as described on figures legends. RT-qPCR technique was performed for all experiments as follows. Cryptococcal cells were washed once with PBS, and then frozen in liquid nitrogen and lyophilized. Cell lysis was performed by vortexing the tubes with the dry pelleted cells using acid-washed glass beads (Sigma Aldrich Co., St. Louis, MO, USA). Three independent sets of RNA samples for each strain were prepared using TRIzol reagent (Invitrogen, Carlsbad, CA, USA), according to the manufacturer’s protocol. Next, RNA samples were treated with DNAse (Promega, Madison, WA, USA), and a total of 300 ng treated-RNA was used for reverse transcription reaction with ImProm-II Reverse transcriptase (Promega, Madison, WA, USA). The RT-qPCR was assessed on a Real-time PCR StepOne Real-Time PCR System (Applied Biosystems, Foster City, CA, USA). PCR thermal cycling conditions had an initial step at 94 °C for 5 min, followed by 40 cycles at 94 °C for 30 s, 60 °C for 30 s, and 72 °C for 60 s. Platinum SYBR green qPCR Supermix (Invitrogen, Carlsbad, CA, USA) was used as reaction mix, with 1 μL of the cDNA (16 ng) template, in a final volume of 20 μL. Each cDNA sample was done in technical triplicates. Melting curve analysis was performed at the end of the reaction to confirm a single PCR product. Thereon, the data were normalized to the actin cDNA levels. Relative expression was determined by the 2^−Δ*CT*^ method (68).

## Ethics statement

The animals were obtained at the Animal Resource Centre, Floreat Park, Western Australia, Australia. The *in vivo* procedures were performed under the protocol number 4254, approved by Western Sydney Local Health District Animal Ethics Committee, accomplished according to the current guidelines of The National Health and Medical Research Council of Australia. The Animal Use Ethics Committee (CEUA/UFRGS) approved the animal experimentation under reference 22488.

## Acknowledgements

We are grateful to Leonardo Nimrichter, Kildare Miranda and Augusto Schrank for helpful discussions. We also thank Hiten Madhani for providing the *ncs1Δ, cna1Δ and cam1Δ* mutant strains, obtained from the Madhani knockout library (http://www.fgsc.net/crypto/crypto.htm), which was created using NIH funding (R01AI100272). We thank Arturo Casadevall for providing mAb 18B7.

## Authors Contribution

Conceived and designed the experiments: E.D.S; C.C.S.; J.T.D.; L.K.

Performed the experiments: E.D.S; J.C.V.R; S.L; H.M; J.S; K.F; D.D.

Analyzed the data: E.D.S; S.L; M.H.V; C.C.S; J.T.D; L.K.

Contributed reagents and materials: M.H.V; C.C.S; J.T.D; L.K.

Wrote the paper: E.D.S; J.T.D; L.K.

---

**Supplementary Figure 1.**
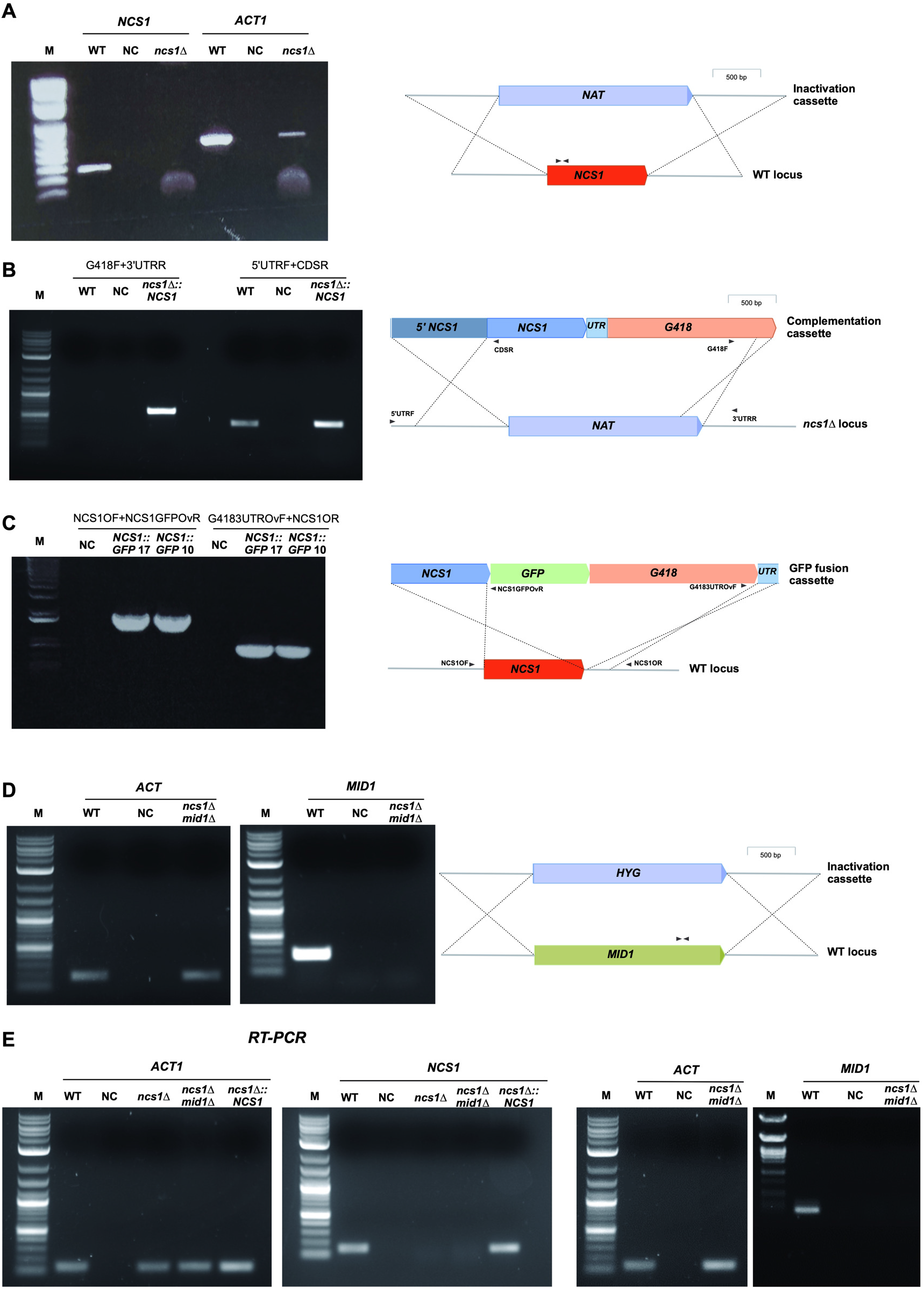
Confirmation of mutant genotypes. (A) The corrected integration of inactivation cassette to generate *ncs1Δ* null strain was evaluated employing PCR with a primer pair that amplify a portion inside the coding region. As loading control, a fragment of the ACT gene was amplified. Left panel: confirmatory PCR analysis. NC, negative control. Right panel, diagram representing the double cross-over at WT *NCS1* locus. (B) Integration of the complementation cassette into the inactivated *ncs1Δ* locus was evaluated with PCR using primers that hybridize inside the complementation cassette (CDSR and G418F primers) and at chromosomal sites outside the double recombination location (5’UTRF and 3’UTRR primers). Each primer pair was used independently to evaluate correct integration at NCS1 CDS upstream site (5’UTRF and CDSR primers), as well as NCS1 CDS downstream site (G418F and 3’UTRR primers). Left panel: confirmatory PCR analysis. NC, negative control. Right panel, diagram representing the double cross-over at *ncs1Δ* locus. (C) Evaluation of the correct integration of *NCS1::GFP* cassette into the WT *NCS1* locus was performed using two primer pairs (NCS1OF + NCS1GFPovR and G4183UTRovF + NCS1OR) independently to assess correct integration at *NCS1* CDS upstream site as well as NCS1 CDS downstream site, respectively. Left panel: confirmatory PCR analysis. NC, negative control. Right panel, diagram representing the double cross-over at WT *NCS1* locus. (D) Evaluation of the correct integration of the inactivation cassette of *MID1* gene in the *ncs1Δ* strain was performed using PCR with a primer pair that amplify a region inside the coding region. As loading control, a fragment of the ACT gene was amplified. Left panel: confirmatory PCR analysis. NC, negative control. Right panel, diagram representing the double cross-over at *MID1* locus. (E) Evaluation of transcript levels of *NCS1* (left gels) or *MID1* (right gels) in distinct mutants and in the WT strain was conducted using RT-PCR with RNA isolated from yeast strains grown in YPD for 24 h. As loading control, the transcript levels of *ACT1* gene were also evaluated. NC, negative control.

**Supplementary Figure 2.**
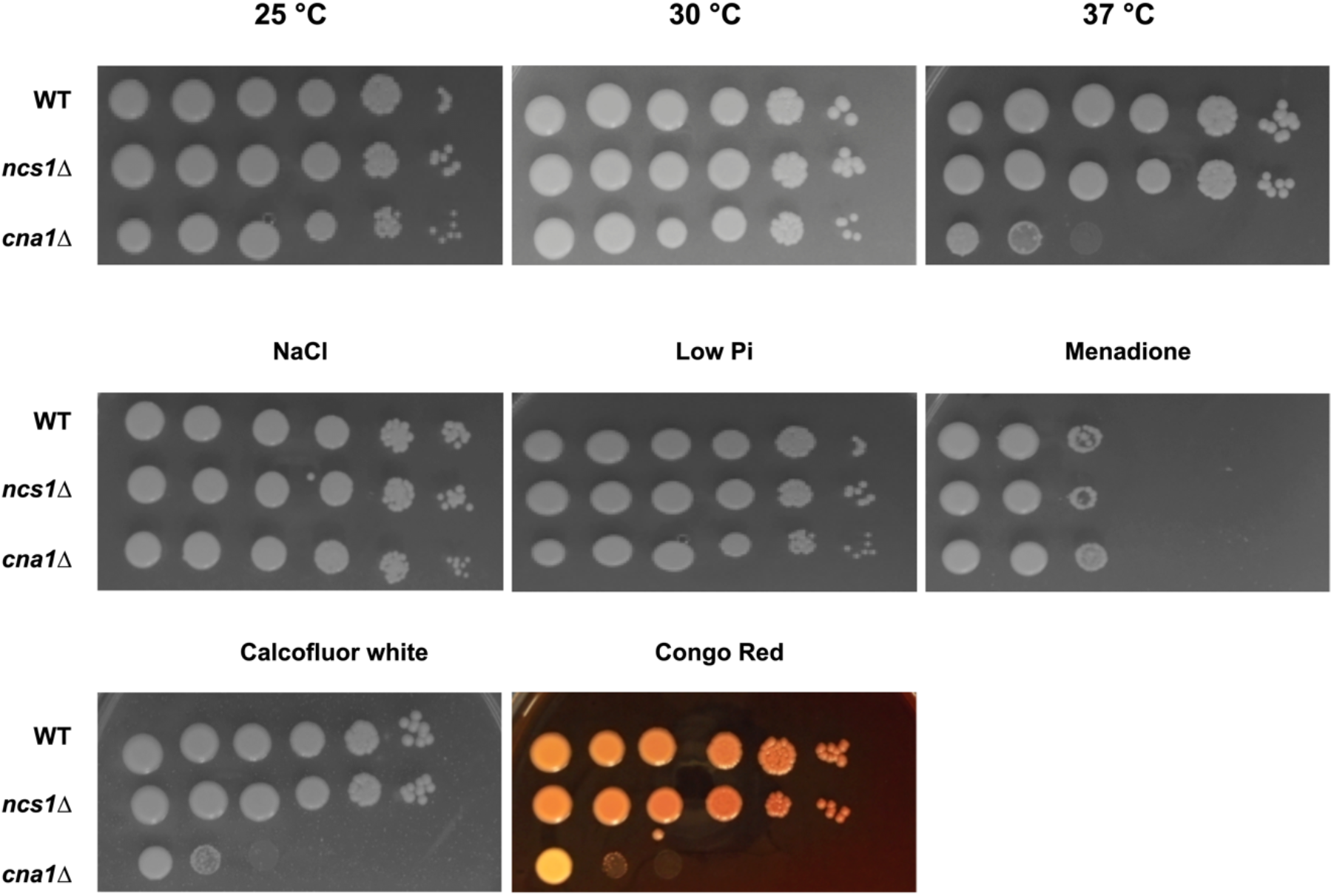
Phenotypic analysis of the *ncs1Δ* null mutant strain. The indicated strains were evaluated by spot plate assay under different stress conditions: altered temperature, saline stress (NaCl 1M), low phosphate and oxidative (menadione) stresses, and cell wall (Calcofluor white and Congo red) stress. Pictures were taken after 48 h of incubation. The calcineurin mutant, *cna1Δ*, was included to assess the degree of phenotypic overlap with *ncs1Δ*.

**Supplementary Figure 3.**
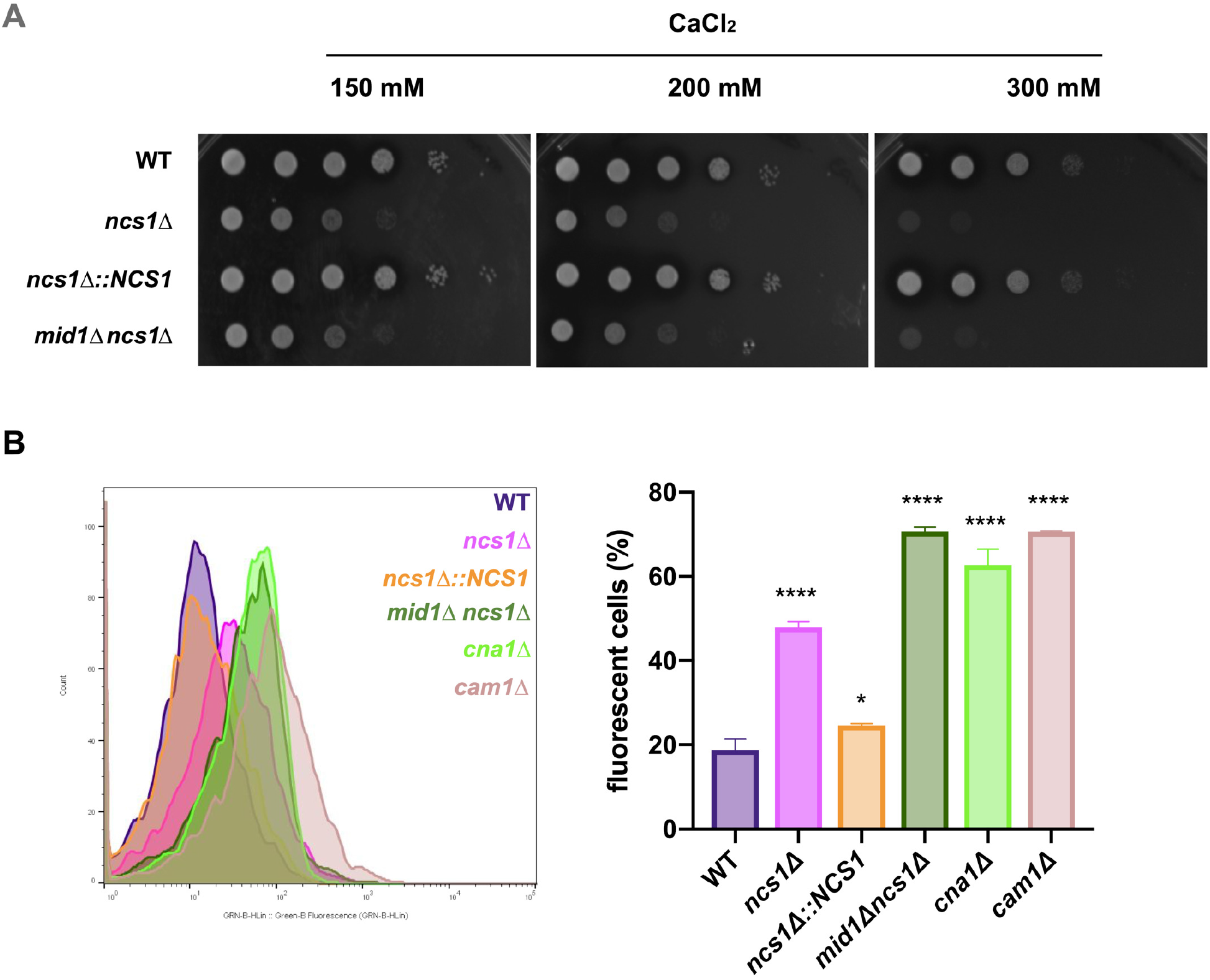
Disruption of *MID1* does not rescue the *ncs1Δ* calcium sensitive phenotype: (A) Spot dilution assay was performed for WT, *ncs1Δ* and *mid1Δncs1Δ* strains in the presence of increasing CaCl_2_ concentrations. Pictures were taken after 48 hours of incubation at 30 °C. (B) The basal level of free intracellular Ca^2+^ in WT, *ncs1Δ, ncs1Δ::NCS1, mid1Δncs1Δ, cna1Δ, cam1Δ* was quantified by flow cytometry following staining with the calcium specific dye, Fluo-4-AM. Left panel represents the histogram of Fluo-4-AM emitted fluorescence of the strains cultivated in YPD medium at 30 °C. Right panel represents the percentage of gated fluorescent cells ± standard deviation (three biological replicates). Mean values were compared using one-way-ANOVA and Dunnet’s as a *post-hoc* for statistical analysis. Statistical significance is represented as **** *p*< 0.0001 and * *p*< 0.05.

**Supplementary Movie 1**. Time lapse video microscopy demonstrating WT bud emergence profile.

**Supplementary Movie 2**. Time lapse video microscopy demonstrating *ncs1Δ* bud emergence profile.

